# Dietary Interventions Modulate Cell Competition and Locomotor Decline in a Alzheimer’s Disease *Drosophila* Model

**DOI:** 10.1101/2025.05.12.653450

**Authors:** Carolina Costa-Rodrigues, Jovin R Jacobs, Joana Couceiro, Catarina Brás-Pereira, Eduardo Moreno

**Author notes:** Correspondence (C-R, C); (B-P, C); (E.M.).

## Abstract

Alzheimer’s Disease (AD) is a neurodegenerative disorder characterised by the accumulation of Amyloid-beta 42 (Aβ42) plaques and cognitive decline. Using *Drosophila* models, we investigated the impact of diet on cell competition, a process that eliminates unfit cells and is involved in AD progression. Cell competition is present in AD flies, where unfit neurons express *fwe*^*LoseB*^ and *azot*, leading to their elimination and motor improvements. In this study, we fed AD and control (healthy) flies with either a yeast-based diet (YBD) or a synthetic diet (SAA) for up to 28 days and evaluated cell competition at the cellular level and locomotion as behavioural output. AD flies fed with YBD exhibited a peak of cell competition at day 14 of feeding, being cyclical over time and followed by a progressive locomotion decline. In contrast, the SAA diet delayed the activation of cell competition until day 21, coinciding with locomotion restoration. Moreover, the synthetic diet postponed Aβ formation, suggesting a slower progression of AD. Overexpression of the *human Flower (hFWE)* isoforms in *Drosophila* photoreceptors revealed their conserved function in regulating cell competition, with *hFWE1* identified as the sole loser isoform in neuronal context. Conversely, *hFWE2*, a winner isoform, induced higher levels of cell competition in a diet-dependent manner. *hFWE3* and *hFWE4* induced a milder effect on cell competition but also have a winner-like function. In YBD-fed AD flies, *hFWE2* failed to promote efficient elimination of unfit cells over time, as these flies exhibited even worse locomotion decline than control flies without hFWE2. In contrast with SAA, expression of hFWE2 induced better locomotion outcomes. Our data seems to indicate that diet may regulate AD progression through cell competition regulation, highlighting the complex interplay between diet, cell competition, and AD progression and providing insights into potential therapeutic strategies.

## Introduction

The prevalence of neurodegenerative diseases is rising significantly among the elderly. Dementia impacts over 55.2 million people worldwide and projections show that will grow up to 78 million by 2030, according to the world Alzheimer Report 2021 (Gauthier S, Rosa-Neto P, Morais JA 2021). Alzheimer’s Disease (AD) has multifactorial and complex nature, which involves neuronal death, extracellular accumulation of Amyloid-β (Aβ), mainly the Aβ42 peptide aggregates, and impaired insulin signalling (Bedse et al. 2015; Selkoe and Hardy 2016; De Strooper and Karran 2016). Patients experience cognitive decline, memory loss, and behavioural impairments due to loss of neuronal processes, and aberrant network activity (Huang and Mucke 2012; Gómez-Isla et al. 1996).

Neurodegenerative diseases (NDDs) are complex due to human intricacy and several hypotheses have been proposed as the cause of AD, yet a unified theory remains to be elucidated (Zhang et al. 2024). *Drosophila melanogaster* has emerged as a key model for NDDs research (Jeon et al. 2020), due to its specific characteristic such as a short life cycle, ample progeny and conservation of fundamental cellular processes and signalling pathways, including nutrient-sensing pathways (Mcgurk, Berson, and Bonini 2015; Ambegaokar, Roy, and Jackson 2010; Costa-Rodrigues, Couceiro, and Moreno 2021). Flies also possess a simpler nervous system compared to humans like blood-brain barrier and are composed by the same cell types, including neurons and glia (Ambegaokar, Roy, and Jackson 2010; Mcgurk, Berson, and Bonini 2015). They can also execute complex motor behaviours and execute memory and learning assays (Jeibmann and Paulus 2009; Coelho et al. 2018a). These characteristics make *Drosophila* a valuable model organism for studying NDDs.

Despite extensive research, AD treatments remain ineffective and often focus on symptoms rather than halting disease progression (Palimariciuc et al. 2023). Recent studies link cell competition to AD, showing that removing unfit neurons restores locomotion in flies(Coelho et al. 2018a). Cell fitness is both relative and context-dependent: it varies according to neighbouring cells, and a fit cell in a context, might be unfit in another (Khandekar and Ellis 2024; Merino, Levayer, and Moreno 2016). When differences in fitness emerge within the tissue, the cell competition mechanism ensures the removal of less fit cells, thereby maintaining tissue and organismal homeostasis, in both invertebrates and vertebrates(Ana Lima et al. 2022; Baker 2020). On a cellular level, the fitter winner cells outcompete the unfit loser cells and dysfunctions in the mechanism is associated with AD and cancer (Esteban-Martínez and Torres 2021; Coelho et al. 2018a; Madan, Gogna, and Moreno 2018; Cong and Cagan 2024).

Tissues rely on different mechanisms to eliminate unfit cells (Costa, Brás-Pereira, and Moreno 2020; Clavería and Torres 2016). In AD context, our laboratory described the role of Fitness-fingerprint-mediated cell competition (Coelho et al. 2018a). Flower (Fwe) isoforms are localized at the membrane, with the fwe^Ubi^ isoform expressed in fitter, winner cells, while the *fwe*^*LoseA/B*^ isoforms are expressed in less fit, loser cells(Coelho et al. 2018a; Portela et al. 2010; Rhiner et al. 2010; Merino, Levayer, and Moreno 2016). The Fwe code is cell-type specific and *fwe*^*LoseA/B*^ triggers cell elimination in epithelia, whereas in neuronal cells only *fwe*^*LoseB*^ is sufficient to induce cell elimination (Merino et al. 2013). The activation of the fitness sensor, *ahuizotl* (*azot)*, occurs downstream of *fwe*^*Lose*^, being required for apoptosis induction through the activation of the pro-apoptotic gene *hid* (Merino et al. 2015b). Loss of *azot* impairs cell competition, reducing lifespan and tissue regeneration (Merino et al. 2015a; Coelho et al. 2018a). Variations in *azot* expression in the gut also modulate lifespan, confirming its importance in ageing (Merino 2023). Recent studies propose that less than 50% of *fwe*^*Lose*^-positive cells express *azot* and die, indicating that these cells can be eliminated in the absence of *azot* (Marques-reis, Hauert, and Moreno 2024). Additionally, *azot*-positive cells can persist in the tissue without triggering apoptosis, which suggests the existence of additional checkpoints before cell elimination (Marquesreis, Hauert, and Moreno 2024). Despite our limited knowledge of the *fwe/azot* pathway, the *Drosophila* homolog of SPARC/Osteonectin family, Sparc, is known to be upregulated in loser cells, protecting these cells from elimination in a cell competition-specific manner, and counteracting the effect of Flower (Portela et al. 2010). Furthermore, researchers showed that Sparc and *Fwe* pathways are independent and act in parallel, being *azot* responsible to integrate the signal of both pathways (Portela et al. 2010).

Although several AD models exist, in mice and flies (reviewed in Costa-Rodrigues, Couceiro, and Moreno 2021), we performed our study using a model characterized by the overexpression of two copies of human Amyloid-β 1–42 (2x hAβ42) carrying a secretion signal peptide (UAS-2x hAβ42), using the Gal4/UAS system (Casas-Tinto et al. 2011). This approach mimics Amyloid Precursor protein (APP) duplication linked to early-onset familial AD, producing more robust phenotypes (Casas-Tinto et al. 2011). Coelho and colleagues demonstrated that in *Drosophila*, the neurons near the hAβ42 plaques express *flower*^*LoseB*^, marking them for elimination through *azot* expression (Coelho et al. 2018a). Furthermore, flies with hAβ42 plaques had motor impairments similar to AD patients, and elimination of these loser neurons was sufficient to restore motor coordination and memory formation (Coelho et al. 2018a). Moreover, downregulating *azot* intensified locomotion impairments, while an extra copy of *azot* enhanced competition and improved motor behaviour. These findings suggest that *azot* activation may help counteract AD-related motor decline(Coelho et al. 2018a).

Recent developments have highlighted dietary and lifestyle changes as an approach to slow AD progression and cognitive decline (Ellouze et al. 2023). Research shows that a Mediterranean diet and diets improving metabolic syndrome phenotypes are key strategies to tackle neurodegenerative diseases (Bianchi, Herrera, and Laura 2021). Diet’s influence on disease outcomes is well established, and some reports show this effect can occur through cell competition. For instance high-fat diet (HFD) decreases apical elimination of Ras^V12^-transformed cells from mice intestinal and pancreatic epithelia, impairing cell competition(Sasaki et al. 2018). High-sugar diet (HSD) promotes tumour growth and metastasis in *Drosophila* by transforming Ras/Src-induced tissue growth into aggressive and metastatic tumours, with cells escaping cell competition (Hirabayashi, Baranski, and Cagan 2013). Hamann et al. (2017) showed that glucose withdrawal induces entosis allowing winner cells to obtain nutrients (Hamann et al. 2017). Additionally, mTOR signalling, was shown to act as a fitness sensor, as unfit cells exhibited decreased in mTOR activity, indicating the involvement of nutrient-sensing pathways in cell competition (Bowling et al. 2018). Hyperinsulinemia promoted tumour growth, allowing cells to escape elimination due to increased protein synthesis (Sanaki et al. 2020). Taken together, these findings open up new possibilities for potential dietary interventions in diseases involving cell competition mechanisms.

Therefore, we questioned how diet influences *azot*-dependent cell competition mechanisms and its implications in Alzheimer’s Disease. Our results show that diet modulates *azot*-dependent cell competition mechanisms in AD flies. Activation of cell competition at 21 days translated into restored locomotion when flies were fed with a synthetic diet (SAA), but not in flies fed with a yeast-based diet (YBD). SAA diet delayed AD progression by delaying the rise of hAβ42 levels, allowing flies to be healthier for longer periods. In contrast, when AD flies were fed with YBD, cell competition was induced earlier at 14 days, as previously shown by Coelho et a. (2018), but flies experienced a gradual locomotion decline, possibly from impairments in unfit cell elimination downstream of *azot*. Therefore, SAA seems to be more beneficial than the YBD for AD flies. Our data suggests that diet can regulate the AD onset by delaying the accumulation of toxic aggregates and promoting elimination of unfit cells. Thus, it provides a foundation for identifying therapeutic approaches to manipulate Fitness-Fingerprint-mediated cell competition to treat age-related diseases.

## Results

### Yeast-based diet hinder cell competition in AD, leading to locomotion decline

Our group showed that at 14 days old, AD flies have more *azot* expression and cell death than control (which we refer to as healthy flies in this work) flies (Coelho et al. 2018a). As AD progresses with age, we decided to extend this study by including older flies. Furthermore, we inquired whether diet affects cell competition and AD progression. Thus, to address how diet modulates *azot*-dependent cell competition and its behavioural implications in AD, we fed healthy flies (*GMR>GFP;azot::mCherry)* and AD (*GMR>hAβ42;azot::mCherry)* flies for 3, 14, 21 and 28 days either on Yeast-Based diet (YBD) or Synthetic (SAA) diet, and evaluate *azot* expression, cell death, elimination of unfit cells and locomotion. We assessed unfit cells by measuring the area of *azot* expression with an *azot::*mcherry reporter (Coelho et al. 2018a) and cell death by the area of TUNEL staining. To evaluate the elimination of unfit cells, we quantified the colocalisation of *azot* with TUNEL staining (which we designated as *azot*-TUNEL-positive cells), using a combination of machine learning and custom written software, as described in Methods section.

Healthy flies fed with a YBD (Fig A-B’, I-I’’’) had drastically increased *azot* expression (Fig1I) by day 21 of feeding which remained constant until day 28. A peak in TUNEL area was detected at day 14 (Fig1 I’), followed by a gradual decline. The area of *azot*-TUNEL positive cells (Fig1I’’) exhibited a similar pattern to *azot* expression over time, with a significant increase by day 21 which plateaued after. These results show an increase in the levels of unfit cells and the rate at which these cells die, potentially correlating with ageing. Furthermore, up to day 28 of feeding, there is a capacity to eliminate unfit cells (Fig 1I’’), with flies maintaining locomotion, which is measured through the activity parameter (Fig1I’’’, FigS1A-A’’’). The previous work from our group, shows that cell competition is associated with improvement in learning and motor function in AD flies(Coelho et al. 2018a). Thus, our findings suggest that this effect occurs also in flies without AD over time, where the increase in unfit cell elimination was sufficient to prevent a drastic locomotion decline in HEALTHY flies.

**Figure 1:**
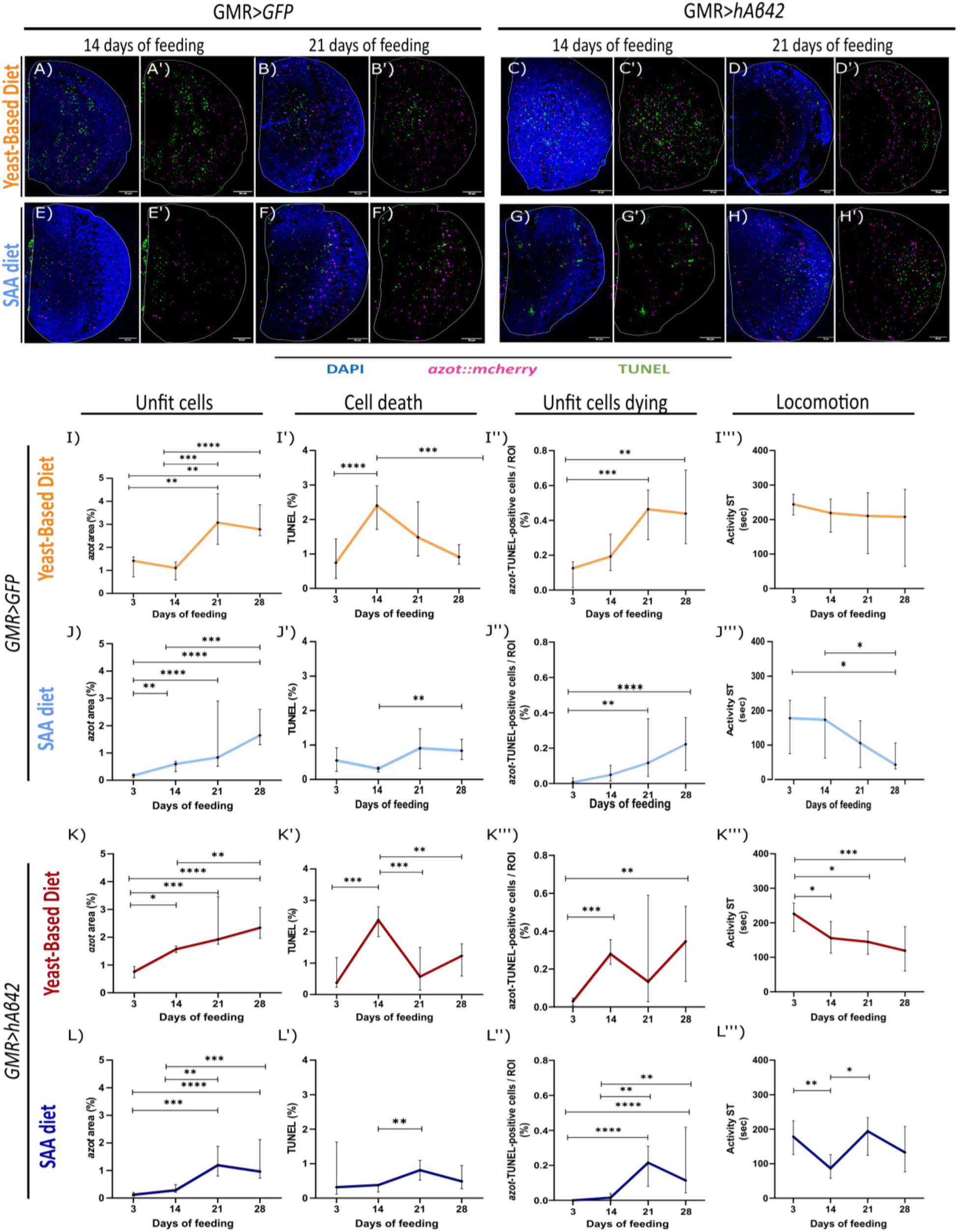
Yeast-Based Diet (YBD) and Synthetic diet (SAA) have different impacts on cell competition. Representative images of optic lobes (OL) of adult healthy flies(A-D’) and AD flies (E-H’) fed for 14 days with YBD (A-B’) and SAA diet (E-F’) and for 21 days with YBD (C-D’) and SAA diet (G-H’). Both healthy and AD flies carry an UAS-LacZ. (A-H’) DAPI for nuclei label (blue), *azot* reporter (magenta) and TUNEL for cell death label (green). Quantification of *azot* expression shows the levels of unfit cells (I and J for healthy flies; K and L for AD flies). Quantification of TUNEL shows the levels of cell death (I’ and J’ for healthy flies; K’ and L’ for AD flies). Quantification of *azot*-TUNEL-positive cells shows the levels of unfit cells dying and is measured by the area of *azot* and TUNEL colocalising (I’’ and J’’ for healthy flies; K’’ and L’’ for AD flies). All quantifications measure the area of signal divided by the region of interest (ROI), which is the optic lobe (%). Quantification of Activity Speed Threshold represents a measure for a fly’s locomotion and is the amount of time that fly’s velocity is above a particular threshold - 2.7 mm/s (I’’’ and J’’’ for healthy flies; K’’’ and L’’’ for AD flies). All flies were fed for 3, 14, 21, and 28 days. For cellular studies (*azot* area, TUNEL area and area of *azot*-TUNEL-positive cells colocalized) n shows the number of optic lobes analysed: (I-I’’) 3d n=13, 14d n=15, 21d n=9 and 28d n=13; (J-J’’) 3d n=18, 14d n=25, 21d n=10 and 28d n=14; (K-K’’) 3d n=6, 14d n=35, 21d n=7 and 28d n=12; and, (L-L’’) 3d n=11, 14d n=30, 21d n=12 and 28d n=16. n shows the number of flies analysed for locomotion analysis: (I’’’’) 3d n=16, 14d n=13, 21d n=9 and 28d n=4; (J’’’) 3d n=24, 14d n=22, 21d n=22 and 28d n=13; (K’’’’) 3d n=23, 14d n=25, 21d n=20 and 28d n=16; and, (L’’’’) 3d n=25, 14d n=19, 21d n=19 and 28d n=20.

In AD flies (*GMR>hAβ42;azot::mCherry)* fed with the YBD (Fig1C-D’, K-K’’’), we observed a significant increase in *azot* expression (Fig1K) at day 14 of feeding, which continuously increased over time. At the same time, TUNEL (Fig1K’), exhibited oscillatory levels over time. It peaks at day 14 of feeding, followed by a decrease on day 21 and a tendency to start rising again on day 28, although it was not statistically significant. A similar pattern was obtained for the *azot*-TUNEL-positive cells (Fig1K’’), with a significant increase by days 14 and 28 compared to day 3 of feeding. AD flies exhibited a gradual locomotion decline (Fig1K’’’, FigS1C-C’’’) despite activation of cell competition on day 14. As previously reported by Coelho et al. (2018), our findings show that YBD promote a peak in cell competition on day 14 in AD. However, over time, the variations in cell death and unfit cell elimination levels might correlate with the decline in locomotion. The fact that there is a continuous rise in *azot* expression, which does not correspond to the pattern of *azot*-TUNEL-positive cells and TUNEL, suggests an accumulation of unfit cells, which might be contributing to the deterioration of locomotor function. These results hint that unfit cell elimination (*azot*-TUNEL-positive cells) is blocked downstream of *azot* activation in flies fed with YBD, and that this diet supports AD progression by worsening mobility. Then, we questioned these cellular and behavioural effects could be differently regulated by a distinct diet.

### SAA Diet Delays Cell Competition and Restores Locomotion in AD Flies

To assess the dietary effects of nutrients on cell competition in AD flies, we fed flies with SAA diet, an exome-matched diet where each nutrient is added separately and in exact concentrations, allowing precise nutrient manipulation (Piper et al. 2017). Flies fed with this diet are phenotypically similar to flies fed with YBD, making it suitable for our studies (Piper et al. 2017). To link changes in *azot*-dependent cell elimination and locomotion to the diet, we replicated the previous experiments in YBD using the SAA diet.

In healthy flies on an SAA diet (Fig1E-F’, J-J’’’), *azot* expression (Fig1 J) increased on day 14. TUNEL levels (Fig1 J’) remained low, increasing from day 21 onwards, being significant only on day 28. After 21 days, flies exhibited a significant increase in *azot*-TUNEL-positive cells (Fig1J’’), which continued to increase on day 28. Locomotor activity (Fig1J’’’, FigS1 F-F’’’) remained constant until day 14, declining afterwards and reaching significantly lower levels on day 28 compared to days 3 and 14. Our results show that despite the rise in unfit cells and unfit cell elimination (*azot*-TUNEL-positive cells) over time, locomotion is not restored, which may correlate with the lack of a peak in cell death at 14 days in SAA-fed flies. Compared with healthy flies fed the YBD (FigS1 E-E’’’), SAA diet induced overall lower levels of *azot* expression, being significant at days 3 and 21 (Fig1I-J, FigS1 E). Regarding TUNEL levels (Fig1I’-J’, S1 E’), SAA diet prevented the peak seen at 14 days with YBD, but by days 21 and 28, both diets induced similar levels. *azot*-TUNEL-positive cells (Fig1I’’-J’’, S1 E’’) were overall lower with the SAA diet, but statistically significant only on day 14. Locomotion (Fig1I’’’-J’’’, FigS1 E’’’) revealed statistical differences on day 3, where YBD-fed flies were more active. Nonetheless, distinct patterns were evident, with SAA-fed flies performing worse than those on YBD. Thus, the YBD diet might be more beneficial for healthy flies compared to the SAA diet, as it allows a peak of cell death at 14 days and higher levels of *azot*-TUNEL-positive cells and *azot* expression, preventing a severe locomotion decline. The SAA diet is optimised for development and egg-laying, but this comes at the expense of lifespan(Piper et al. 2017). As a result, we cannot exclude that the decline in locomotion seen on the SAA diet may also be age-related rather than driven by cell competition.

When AD flies were fed an SAA diet (Fig1G-H’, L-L’’’), we observed a spike in the number of *azot*-expressing cells (Fig1L), TUNEL (Fig1L’) and *azot*-TUNEL-positive cells (Fig1L’’) on day 21, suggesting that cell competition was induced after 21 days of feeding. AD flies’ locomotion (Fig1L’’’, FigS1H-H’’’) tracked the cellular results by being restored also at day 21 of feeding. These results suggest that the SAA diet allows activation of cell competition by day 21 which correlates with recovery of locomotion. A comparison between diets (FigS1J-J’’’) showed that SAA diet led to overall lower *azot* expression than YBD (Fig1C-D’, G-H’, K-L, FigS1 J), being significantly lower on days 14 and 28. Furthermore, TUNEL levels (Fig1K’-L’, FigS1J’) were also different, with SAA abolishing the peak seen at day 14 on flies fed with YBD. In terms of *azot*-TUNEL-positive cells (Fig1K’’-L’’, FigS1J’’), AD flies on YBD showed a peak at day 14, while SAA-fed flies, shifted to day 21. These results indicate that feeding AD flies with an SAA diet delayed the activation of cell competition to day 21. Also, despite the variations in locomotion (Fig1K’’’-L’’’, FigS1J’’’) between YBD and SAA diet were not statistically significant, there was a noticeable tendency. SAA-fed flies restored locomotion on day 21, whereas YBD-fed flies exhibited a gradual decline over time. Altogether, results show that the YBD and SAA diets have different impacts on *azot*-dependent cell competition modulating the fly’s locomotion. Moreover, YBD diet induced more and earlier fitness differences than SAA diet. We then wondered if the effect seen with SAA diet correlates with the levels of hAβ42.

### SAA delays accumulation of hAβ42

We hypothesise that the SAA diet may slow down locomotion decline by limiting hAβ42 accumulation, which may restrain toxicity and the increase in unfit cells, postponing cell competition activation. We fed AD flies with YBD and SAA diets for 3, 14, and 21 days and evaluated the levels of hAβ42 using a specific antibody and measuring the area of signal (Fig2A-B’, C). We simultaneously assessed *azot* expression (Fig2A-B’, D). YBD-fed flies increased hAβ42 levels by day 14, which kept higher on day 21 compared to day 3 (Fig2C), coinciding with the rise in *azot* expression on day 14 (Fig2D) and confirming our previous *azot* results (Fig1K). Compared to YBD, SAA-fed flies exhibited approximately half the hAβ42 levels and *azot* signal at day 14 (Fig 2 B-B’, D-E). However, by day 21, SAA and YBD-fed flies had comparable levels of hAβ42 (Fig2C), but SAA-fed flies exhibited lower levels of *azot* expression (Fig2D), despite being higher than those at day 14 on the SAA diet. Together, results suggest that the SAA diet delays the accumulation of hAβ42 protein and *azot*-expressing cells, postponing cell competition to day 21 (FigS1J-J’’’). The lower amount of hAβ42 in SAA-fed flies might indicate a delayed AD onset, which could explain locomotion restoration at day 21. In contrast, locomotion on YBD-fed flies gradually declines (Fig2E). While we have identified how the SAA diet may be impacting cell competition in AD, we remain uncertain about the cause of the gradual decline in locomotion observed in flies fed with YBD, despite the activation of cell competition at 14 days of feeding.

**Figure 2:**
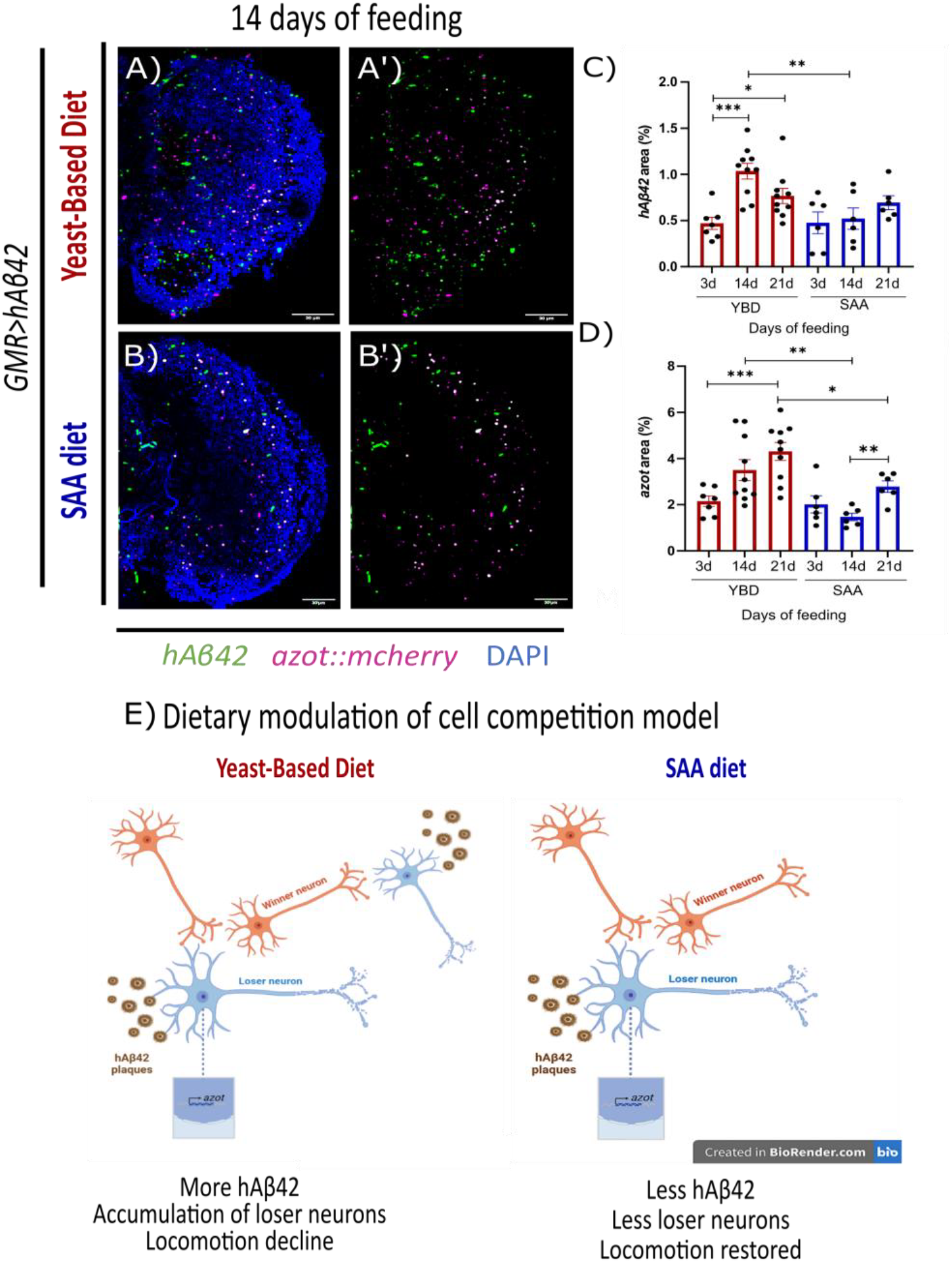
SAA diet prevents high levels of hAβ42 compared to Yeast-Based Diet (YBD). Representative images from the optic lobe of AD flies fed with YBD (A-A’) and SAA diet (B-B’) for 14 days, with DAPI (blue), *azo*t::mcherry (magenta) and hAβ42 (green). (C) Quantification of hAβ42 area (%) normalized to the area of optic lobe. (D) Quantification of azot area (%) normalized to the area of optic lobe. (C-D) Flies were fed for 3, 14 and 21 days. Redd bars represent YBD and blue bars represent SAA diet. (E) Working Model – Feeding YBD led to higher levels of hAβ42, leading to more loser neurons (blue) and locomotion decline. While SAA induced lower levels of hAβ42, less loser neurons and maintenance of winner neurons (orange), allowing restoration of locomotion upon cell competition.

### hFWE isoforms are functionally conserved in *Drosophila* adult neuronal tissue

In the *Drosophila* AD model, *hAβ42* is expressed under the GMR domain, secreted to the extracellular space, and forms aggregates that are randomly distributed in the optic lobe (Casas-Tinto et al. 2011). Hence, creating a heterogeneous environment with neurons of varying fitness status. As demonstrated by Coelho et al. (2018), the neurons near hAβ42 plaques exhibit loser traits, including expression of *fwe*^*LoseB*^ and *azot*. In this context, exacerbation of fitness status differences has beneficial effects, restoring flies’ locomotion (Coelho et al. 2018a). To unravel whether the accumulation of *azot*-expressing cells reflects a blockage of cell elimination, we overexpressed a *fwe* winner isoform to promote fitness differences and force cell competition, potentiating unfit cells’ elimination (Coelho et al. 2018a). We reasoned that if the pathway is compromised downstream of *azot*, expressing a winner isoform would activate cell competition and potentially restore locomotion. Previously, our group confirmed the conservation of *fwe* in humans, describing four *human Flower (hFWE)* isoforms, with *hFWE1* and *hFWE3* functioning like loser isoforms, while *hFWE2* and *hFWE4* behave as winners (Madan et al. 2019). Thus, we decided to use an hFWE winner isoform to potentiate cell competition. To do so, we first needed to assess the functional conservation of hFWE isoforms in *Drosophila* by overexpressing individually each isoform in the AD (Fig3A-I) and healthy (Fig3J-R) flies. We proceed by assessing *azot* expression, cell death (TUNEL), and the elimination of unfit cells (*azot*-TUNEL-positive cells) at 14 days. In AD flies, the overexpression of a loser hFWE isoform in a tissue surrounded by loser cells associated to near the hAβ42 plaques, should decrease cell competition due to increased tissue fitness homogeneity, which would be translated into less *azot* and *azot*-TUNEL positive cells. In contrast, if an hFWE isoform acts like a winner isoform, it will increase fitness differences between photoreceptors and surrounding tissue containing the hAβ42 plaques. Thus, potentiating cell competition and having the opposite effect on the amount of *azot* and *azot*-TUNEL positive cells.

Compared to LacZ control (Fig3A-A’), our results show that in AD flies *hFWE1* (Fig3C-C’) functions similarly to *fwe*^*LoseB*^ (Fig3B-B’), hindering competition and functioning as a loser, by decreasing *azot* expression (Fig3G), TUNEL (Fig3H) and *azot*-TUNEL-positive cells (Fig3I). In contrast, overexpression of *hFWE2* (Fig3D-D’) has the opposite effect on cell competition, inducing the increase of *azot* expression (Fig3H) and *azot*-TUNEL-positive cells (Fig3I), thus behaving as a winner isoform. *hFWE3* and *hFWE4* (Fig3E-E’, F-F’) tend to behave like *hFWE2*, suggesting a winner phenotype, although significant differences occur only when compared to loser isoforms (Fig3G-I). In healthy flies (Fig3J-R), *hFWE1* (Fig3L-L’) and *fwe*^*LoseB*^-expressing flies (Fig3K-K’) show no major differences in the *azot*, meaning that in non-sensitized *hAβ42* context, these isoforms are not sufficient to induce cell competition. *hFWE2* significantly increased *azot* expression (Fig3M-M’,P) compared to the controls, suggesting an increase in fitness differences and thus, cell competition. The remaining hFWE isoforms (*hFWE3* and *hFWE4* – Fig3N-N’ and O-O’, respectively) had minimal effects on *azot* expression (Fig3P) and cell death (Fig3Q) in a healthy context.

**Figure 3:**
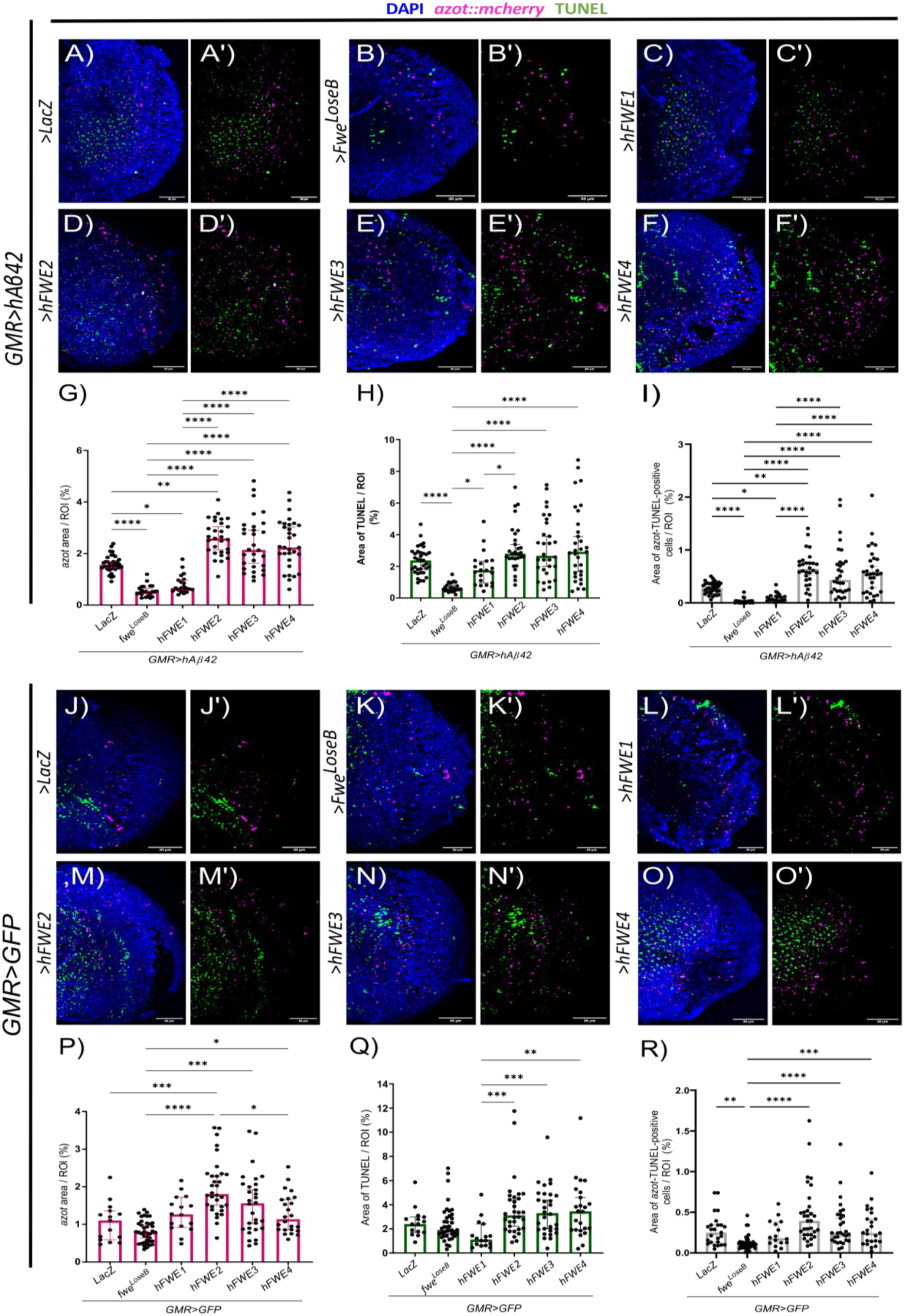
Functional conservation of *hFWE* isoforms in Drosophila optic lobe. Representative images of optic lobes of flies fed with YBD for 14 days and overexpressing *LacZ* (A-A’ in AD, J-J’ in healthy), *fwe*^*LoseB*^ (B-B’ in AD, K-K’ in healthy), *hFWE1* (C-C’ in AD, L-L’ in healthy), *hFWE2* (D-D’ in AD, M-M’ in healthy), *hFWE3* (E-E’ in AD, N-N’ in healthy), *hFWE4* (F-F’ in AD, O-O’ in healthy), showing the signals of interest: DAPI (blue), *azo*t::mcherry (magenta) and hAβ42 (green). (H, P) Quantification of *azot* expression by *azot*::mcherry reporter signal area normalized to ROI (ROI is the optic lobe) in either AD and healthy, respectively (%). (H-Q) Quantification of TUNEL signal area normalized to ROI (%). (I-R) The quantification of *azot*-TUNEL-positive cells is measured by the area of *azot* reporter signal that is colocalized with the TUNEL signal and normalized to the ROI (%). Our n represents the number of optic lobes, in AD flies LacZ n=35; fwe^LoseB^ n=23; hFWE1 n=21; hFWE2 n=29; hFWE3 n=30; hFWE4 n=31; in healthy flies *LacZ* n=15; *fwe*^*LoseB*^ n=39; *hFWE*1 n=14; *hFWE*2 n=32; *hFWE*3 n=30; *hFWE*4 n=24.

Our results reveal that hFWE1 acts as loser isoform, whereas hFWE3, although reported as loser isoform by Madan et al. (2019), does not fulfil this role in neuronal tissue. Thus, mirroring the behaviour of *Drosophila* loser isoforms, where *fwe*^*LoseA*^ does not function as loser in neuronal tissue, while *fwe*^*LoseB*^ function as a loser in both neuronal and epithelial tissues(Merino et al. 2013). These findings demonstrate the functional conservation of *hFWE* isoforms in *Drosophila* and highlight the role of these isoforms in regulating cell competition. We show that *hFWE1* is the only loser isoform in the eye neuronal tissue, similar to *fwe*^*LoseB*^ as previously shown by Rhiner et al (2010), while *hFWE2* acts like a winner isoform, with *hFWE3* and *hFWE4* having a minor impact in cell competition but acting as winner isoforms. After settling the winner-like behaviour of *hFWE2* in *Drosophila*, we used this isoform to potentiate competition and evaluated impairments downstream of *azot* in an attempt to unravel YBD effects.

### Winner isoform *hFWE2* promotes unfit cell elimination in a diet-dependent manner

To assess whether increasing relative fitness differences enhances unfit cell elimination and restores locomotion in flies fed with YBD, we overexpressed *hFWE2* on healthy and AD flies fed with YBD or SAA diets for 14, 21, and 28 days. *hFWE2* expression should improve unfit cell elimination and restore locomotion, if *fwe/azot* pathway is not inhibited downstream of *azot* expression. Again, we evaluated cell competition and locomotion.

In healthy flies fed with YBD and expressing *hFWE2* (Fig4A-C’, dashed line in M-M’’’), *azot* expression (Fig4M) and *azot*-TUNEL-positive cells (Fig4M’’) remained stable over time, while TUNEL gradually decreased (Fig4M’). Flies exhibited a locomotion decline over time (Fig4M’’’). Comparing healthy flies expressing *LacZ* vs *hFWE2* (solid vs dashed line, respectively in Fig4M-M’’’), revealed that *hFWE2*-expressing flies exhibit more cell competition at 14 days, being statistically significant for *azot* expression (Fig4M) and *azot*-TUNEL-positive cells (Fig4M’’). However, *hFWE2* flies exhibited worse locomotion than *LacZ* flies (Fig 4 M’’’). These results indicate in healthy flies *hFWE2* do not restore locomotion despite higher levels of unfit cells; instead, induce an even worse phenotype than *LacZ*. AD flies fed with YBD and expressing *hFWE2* (Fig4G-I’, dashed line in O-O’’’) revealed that *azot* expression (Fig4O) increases, while TUNEL (Fig4O’) and *azot*-TUNEL-positive cells (Fig4O’’) decrease over time. This inverse correlation explains the tendency for *hFWE2*-expressing flies to decrease locomotion compared to *LacZ* flies (Fig4O’’’). Similar to healthy flies, expression of *hFWE2* in AD flies at 14 days, induced more *azot* expression and *azot*-TUNEL-positive cells at 14 days than the *LacZ* (solid vs dashed line, respectively, Fig4O-O’’). *hFWE2* induced fitness differences, however, did not translated into better mobility performance of these flies (Fig4O’’’). One hypothesis for these results is that an unidentified player is preventing the efficient elimination of unfit cells over time, which could explain the TUNEL decrease (Fig4O’). We propose that in AD flies fed with YBD, *hFWE2* expression induces higher fitness differences between photoreceptor neurons and surrounding cells, but unfit cells are not efficiently eliminated from the tissue, promoting their accumulation and locomotion decline, even more than LacZ flies. The idea that, in neuronal tissue, *azot* expression is not sufficient to induce cell elimination is in line with recent studies from our group (Marques-reis, Hauert, and Moreno 2024).

**Figure 4:**
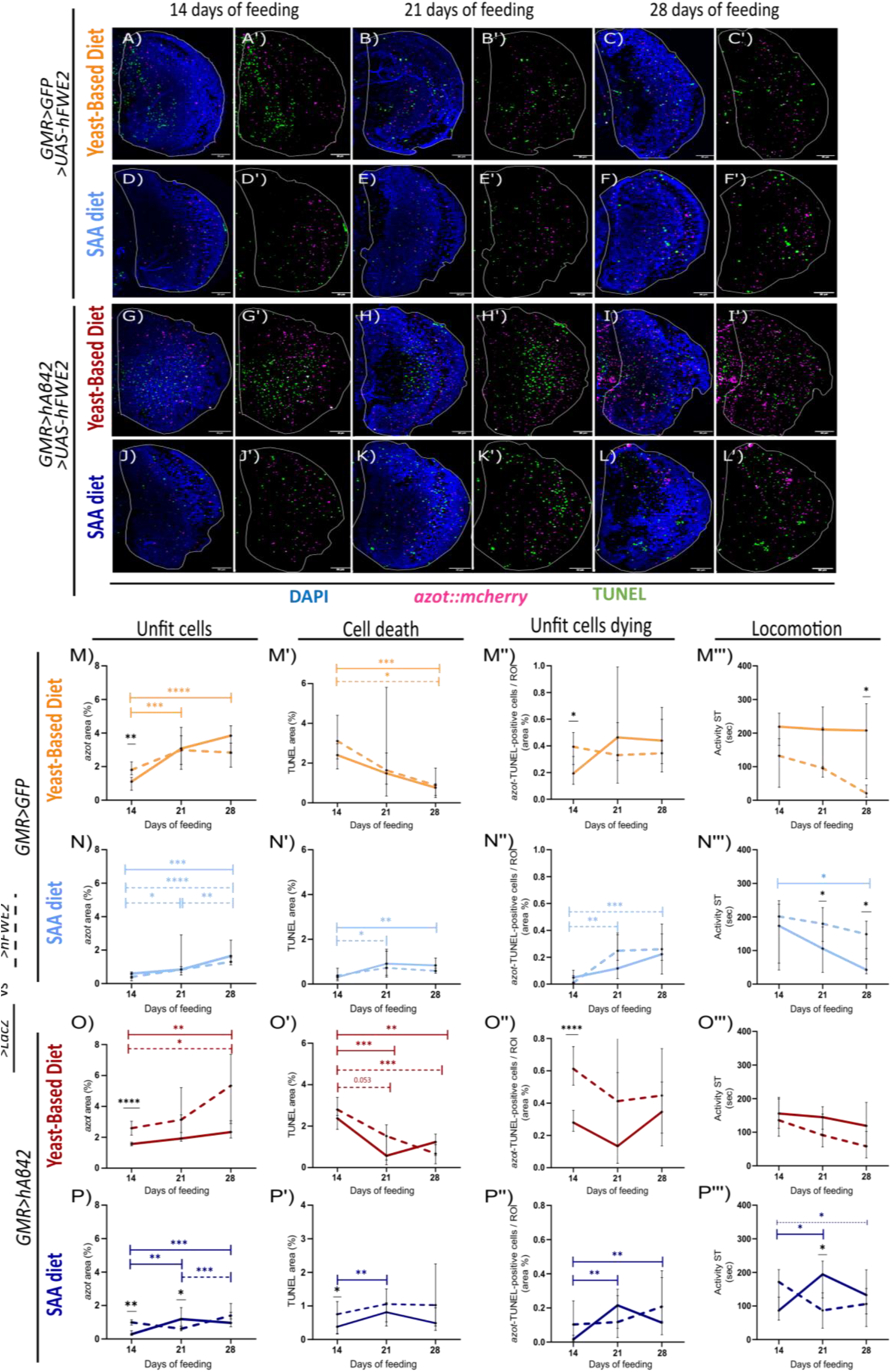
Winner isoform overexpression was not enough to restore locomotion in Yeast-Based Diet. Representative images of optic lobes (OL) of adult healthy flies fed with YBD (A-C’) and with SAA (D-F’); and AD flies fed with YBD (G-I’) and SAA diet (J-L’) fed for 14 to 28 days. (A-L’) DAPI for nuclei label (blue), *azot* reporter (magenta) and TUNEL for cell death label (green). (I-L’’’) Dashed lines represent flies expressing *hFWE2* and solid lines represent *LacZ*-expressing flies (same data from Fig 1). Quantification of *azot* expression shows the levels of unfit cells (I and J for healthy flies; K and L for AD flies). Quantification of TUNEL shows the levels of cell death (I’ and J’ for healthy flies; K’ and L’ for AD flies). Quantification of *azot-*TUNEL-positive cells shows the levels of unfit cells dying and is measured by the area of *azot* and TUNEL colocalising (I’’ and J’’ for healthy flies; K’’ and L’’ for AD flies). All quantifications measure the area of signal divided by the region of interest (ROI), which is the optic lobe (%). Quantification of Activity Speed Threshold represents a measure for a fly’s locomotion and is the amount of time that fly’s velocity is above a particular threshold - 2.7 mm/s (I’’’ and J’’’ for healthy flies; K’’’ and L’’’ for AD flies). All flies were fed for 14, 21, and 28 days. For cellular studies (*azot* area, TUNEL area and area of *azot*-TUNEL-positive cells colocalized) n shows the number of optic lobes analysed: (I-I’’) +*LacZ* at 14d n=15, at 21d n=9 and at 28d n=13; +*hFWE*2 flies14d n=32, at 21d n=6 and at 28d n=9; (J-J’’) +*LacZ* flies at 14d n=25, at 21d n=10 and at 28d n=14; +*hFWE*2 flies14d n=7, at 21d n=21 and at 28d n=30; (K-K’’) +*LacZ* flies at 14d n=15, at 21d n=9 and at 28d n=13; +*hFWE*2 flies14d n=32, at 21d n=6 and at 28d n=9; (L-L’’) +*LacZ* flies at 14d n=25, at 21d n=10 and at 28d n=14; +*hFWE*2 flies14d n=7, at 21d n=21 and at 28d n=30. In locomotion studies, n shows the number of flies analysed: (I’’’) +*LacZ* flies at 14d n=13, at 21d n=9 and at 28d n=4; +*hFWE*2 flies at 14d n=7, at 21d n=12 and at 28d n=6; (J’’’)+*LacZ* flies at 14d n=22, at 21d n=22 and at 28d n=13; +*hFWE*2 flies14d n=7, at 21d n=36 and at 28d n=20; (K’’’)+*LacZ* flies at 14d n=25, at 21d n=20 and at 28d n=16; AD+*hFWE*2 flies at 14d n=39, at 21d n=13 and at 28d n=4; +*LacZ* flies at 14d n=19, at 21d n=19 and at 28d n=20; +*hFWE*2 flies14d n=5, at 21d n=18 and at 28d n=17.

We did the same evaluation in SAA-fed flies and observed that in healthy flies expressing *hFWE2* (Fig4 D-F’, dashed line in N-N’’’ dashed line), *azot* expression (Fig4N) increased continuously over time, while TUNEL levels (Fig4N’) and *azot*-TUNEL-positive cells (Fig4N’’) spiked by day 21 and remained constant by day 28. These flies presented stable levels of locomotion over time (Fig4N’’’), indicating that *hFWE2* induces unfit cells and that elimination of these cells from day 21 onwards (Fig4N’’dashed line), is sufficient to prevent a drastic locomotion decline. Comparing *hFWE2*-expressing flies with control flies expressing *LacZ* revealed no statistically significant differences in any of the cellular parameters evaluated (dashed vs solid line in Fig4N-N’’). However, flies expressing *hFWE2* have better locomotion than the *LacZ* ones at 21 and 28 days (Figs4N’’’). These results suggest that although the cell competition parameters are similar between both conditions, *hFWE2* might improve tissue fitness in SAA diet. Furthermore, expression of *hFWE2* might increase differences in cell fitness, promoting unfit cell elimination and leading to a better locomotion outcome. These results show that in SAA diet the pathway is not hindered downstream of *azot*, as cells are efficiently eliminated.

Analysis of *hFWE2* expression in HEALTHY flies fed SAA diet versus YBD (Fig4, FigS2 A-A’’’) revealed that SAA-fed flies, exhibited less *azot* expression across all time points overtime (Fig S2 A). TUNEL (Fig S2 A’) and *azot*-TUNEL-positive cells (FigS2 A’’), both parameters were significantly lower on SAA-fed flies at day 14. However, over time the levels of TUNEL and *azot*-TUNEL-positive cells are similar in both diets. Moreover, flies fed with an SAA diet had better locomotion over time than those fed with a YBD (FigS2A’’’), suggesting that the lower levels of cell competition in SAA-fed flies might be attributed to a better fitness status of cells, and do not to an impairment of the cell elimination mechanisms.

Regarding AD flies expressing *hFWE2* fed with SAA diet (Fig4J-L’, dashed line in P-P’’’), we observed fluctuating levels of *azot* expression with higher levels on day 28 (Fig4P), while TUNEL (Fig4P’) and *azot*-TUNEL-positive cells (Fig4P’’) levels remained stable over time. Locomotion declines over time (Fig4P’’’), which correlates with the fluctuation in the levels of unfit cells over time. When compared to the control flies expressing *LacZ* (dashed vs solid line in Fig4P-P’’’), expression of *hFWE2* induced higher levels of *azot* expression (Fig4P), TUNEL (Fig4P’) on day 14 of feeding, in contrast with *LacZ-*expressing flies. Regarding the *azot*-TUNEL-positive cells (Fig4P’’), there is a similar effect at 14 days. These results suggest that *hFWE2* expression induced an earlier cell competition onset in AD flies fed with SAA diet. Locomotion also tracked these cellular effects, leading to more active *hFWE2* flies on day 14 (Fig4P’’’). When comparing AD flies fed with YBD and SAA diet (Fig4O-P’’’, FigS2B-B’’’), we observed that YBD-fed flies expressing *hFWE2* showed overall higher *azot* levels (FigS2B), *azot*-TUNEL-positive cells (FigS2B’’). Regarding cell death, YBD-fed flies present higher TUNEL levels initially (FigS2B’), but over time these levels drastically decrease, reaching similar levels to those found in SAA-fed flies on days 21 and 28. However, regardless of the diet, these flies exhibit comparable locomotion declines (FigS2B’’’) regardless of diet. Overall, our data shows that the expression of *hFWE2* in HEALTHY flies led to better locomotion when flies were fed with SAA diet, and in AD flies *hFWE2* failed to significantly improve locomotion in the same diet.

### Foxo and Akt seem to be involved in the regulation of *azot* expression in AD flies

There is a well-established correlation between AD and insulin signalling (IIS). Several studies suggest that AD may be a degenerative metabolic disease being driven by impairments in brain insulin response (Bedse et al. 2015). In AD patients, there is aberrant insulin signalling due to inhibition of the pathway downstream of the Insulin receptor (Bedse et al. 2015). The inhibition of protein kinase B (AKT) prevents FOXO from being phosphorylated, allowing its translocation to the nucleus and consequently target gene activation (Hay 2011). Given the pivotal role of FOXO in metabolism and cell death (Kramer et al. 2003), we hypothesised whether Foxo could regulate *azot* expression. To test our hypothesis, we inhibited *foxo* function either by overexpressing double-stranded RNA (UAS-ds-Foxo) or *akt* (UAS-*akt*), and evaluated the number of *azot*-positive cells in YBD-fed flies at day 14. The results showed that when *foxo* is inhibited either by ds-Foxo (FigS3B,D) or *akt* (FigS3C,D), the number of *azot*-positive cells decreased compared to control flies expressing *LacZ* (FigS3A,D). This result suggests that Foxo regulates *azot* expression at 14 in AD flies fed with YBD, which could be a direct or indirect effect. Moreover, these results offer insight into the potential relationship between IIS, AD and cell competition. IIS may influence cell competition in AD, through the regulation of *azot* expression in a Foxo-dependent manner. Further studies are needed to confirm this mechanism associated with AD.

## Discussion

The prevalence of neurodegenerative diseases such as Alzheimer’s Disease (AD) has increased worldwide, together with life expectancy. However, despite a wider range of studies on the aetiology and risk factors in the last few years, the treatment options focus on symptomatic relief rather than stopping/delaying disease progression (Palimariciuc et al. 2023). Recently, new approaches have emerged, and dietary patterns have been shown to modulate cognitive decline and prevent disease progression (Ellouze et al. 2023). Cell competition is a surveillance mechanism shown to be involved in Alzheimer’s disease, with a beneficial effect (Coelho et al. 2018a). Our findings show that activation of *azot*-dependent cell competition in the AD fly model is diet-dependent (Fig1). We observed a correlation between cell competition activation with efficient elimination of unfit cells and locomotion restoration (Fig1 and Fig4). Our results showed that diet modulates these events. Synthetic diet has the potential to delay unfit cell elimination, with locomotion being restored when cells are efficiently eliminated. Our findings also show that synthetic diet delays AD progression by delaying the accumulation of hAβ42 protein (Fig2).

### Yeast-based diet leads to locomotion decline despite cell competition activation

In AD flies fed a yeast-based diet (YBD) (Fig1K-K’’’), *azot*-dependent cell competition was induced earlier than in those on a synthetic diet (SAA diet) (Fig1L-L’’’). Despite *azot* activation at 14 days, unfit cells continued to rise (Fig1K), while unfit cells dying followed an oscillatory pattern (Fig1K’’), suggesting that some unfit cells persist in the tissue, driving locomotion decline over time (Fig1K’’’). Our results showed that fluctuations in unfit cell elimination strongly correlate with locomotion deterioration and the parameters evaluated must align to restore locomotion. Given prior studies showing that cell competition benefits AD flies and improves locomotion (Coelho et al. 2018a), we hypothesised that YBD may impair *azot*-dependent cell competition downstream of *azot*, allowing unfit cells to persist (Fig2E). This raises the possibility of an unknown factor counteracting *azot* expression cyclically, causing oscillatory effects on unfit cell elimination.

To address impairments downstream of *azot* in YBD-fed AD flies, we overexpressed a *hFWE* winner isoform, thereby intensifying cell competition. However, overexpressing *hFWE2* increased *azot* expression over time while cell death and elimination of unfit cells gradually declined (Fig4O’-O’’), failing to restore locomotion (Fig4O’’’) and suggesting impairments in unfit cell removal. Our study reveals that hFWE isoforms are functionally conserved in *Drosophila* adult neuronal tissues (Fig3), as hFWE1 acts like a loser isoform and hFWE2 acts like a winner isoform. hFWE3 and hFWE4, seem to act as winner isoforms as they are significantly different from loser isoforms and similar to hFWE2. Expressing hFWE alongside *hAβ42* enhances the humanisation of this AD model, improving studies on AD – cell competition link.

Research on YBD-fed flies revealed a cyclical unfit cell accumulation and elimination in *azot*^*-/-*^ flies, suggesting an *azot*-independent mechanism removes suboptimal cells when *Fwe*-mediated cell competition is compromised (Marques-Reis, Hauert, and Moreno 2024). In flies with intact cell competition, less than 50% of loser cells expressing *fwe*^*LoseB*^, activate *azot* and undergo apoptosis, hinting that unfit cells aren’t always eliminated (Marques-reis, Hauert, and Moreno 2024). Researchers proposed that other parallel pathways may counteract the effect of *fwe*^*LoseB*^, challenging *azot*’s role as the ultimate fitness sensor (Marques-reis, Hauert, and Moreno 2024). Different studies suggest that azot integrates signals from multiple factors including relative levels of fwe ^lose/win^; sparc levels, and the percentage of winner neighbouring cells (Merino et al. 2015b; Portela et al. 2010; Levayer, Hauert, and Moreno 2015). *Sparc*, the *Drosophila* homolog of SPARC/Osteonectin family, is upregulated in loser cells, counteracting *fwe* effects and preventing apoptosis during Fwe-independent cell competition(Portela et al. 2010). It is plausible that Sparc expression increases in YBD-fed flies, hampering unfit cell removal downstream of *azot*.

In accordance, high levels of SPARC-like 1 were detected in AD patients’ cerebrospinal fluid, proposing SPARC as a potential biomarker (Vafadar-Isfahani et al. 2012). In mice neural stem cells, Testican2, regulated by Brd4 (Bromodomain Containing 4) and responsible for Sparc degradation, regulates cell competition, ensuring the elimination of unfit cells (Li et al. 2024). Additionally, dementia patients exhibited higher levels of Sparc and low levels of Brd4, and mice carrying Brd4 mutations detected in dementia patients, present impairments in neural stem cell competition (Li et al. 2024). Reduced mRNA levels of winner isoforms m*Fwe*3 and m*Fwe*4 were found in the cortex of *Brd4*KO aged mice, where Testican2 does not degrade Sparc. These findings indicate a decline in cell competition capacity and open the possibility for Sparc to regulate Flower in mammals (Li et al. 2024). Although Portela et al. (2010) stated that *Fwe* and Sparc act in parallel pathways in the epithelial imaginal disc, neuron-specific mechanisms may differ due to the importance of neurons to the fly’s visual system (Portela et al. 2010). This mechanism might work in our AD model since Fs(1)h (*female sterile* (*1*) *homeotic*), the *Drosophila* ortholog of Brd4, reduces cell death in the *Drosophila* AD model (Marques dos Reis et al. 2017). We suggest that variations in endogenous *Sparc* might counteract *azot* in YBD-fed flies since this diet promotes high levels of unfit cells, which would be detrimental to the flies’ visual system due to excessive elimination of photoreceptor neurons. Furthermore, YBD seems to promote more hAβ42 levels than the SAA diet (Fig2D) thus the progression of AD.

Cell functions rely on nutrients provided by the diet. Thus, nutritional composition may influence the balance between the *Fwe* pathway and parallel pathways that prevent unfit cell removal. Cellular stress levels can favour one pathway instead of the other. In humans, insulin, IGF-1, or LEPTIN are SPARC stimulators, showing that nutrient-sensing pathways can modulate SPARC (Kos and Wilding 2010). Thus, our findings revealing that efficient elimination of unfit cells is diet-dependent are supported by previous research. Based on this, we wondered whether some key metabolic players could be modulating *azot* expression. Downregulation of *foxo* or overexpression of *akt* was able to downregulate *azot* levels compared to control AD flies. Both genes are involved in the IIS pathway, which is compromised in AD (Bedse et al. 2015). When IIS is impaired, Akt is unable to phosphorylate Foxo, allowing its nuclear translocation and activation of target genes, such as *azot* (Bedse et al. 2015). Thus, *foxo* downregulation limits *azot* expression. Nevertheless, our results cannot discriminate the nature of this effect, whether it is direct or indirect, but are consistent with the perspective that a cell with impaired energetic metabolism will exhibit a decreased fitness, thereby fostering a competitive phenotype. *foxo* has previously been implicated in cell competition, with *Foxo3* being upregulated in loser cells due to stressful conditions (A. Lima et al. 2020).

### Synthetic diet delays AD progression

AD flies fed with SAA diet (Fig1L-L’’’) exhibit a spike in *azot*, TUNEL and *azot-*TUNEL-positive cells only at 21 days later than those on YBD (Fig1K-K’’’). Locomotion restoration correlates with the increase in these parameters (Fig1L’’’) in SAA-fed flies, while YBD-fed flies show locomotion decline (Fig1K’’’). The SAA diet benefits AD flies by efficiently eliminating unfit cells and restoring locomotion when cell competition is triggered (Fig2E). Thus, SAA seems to delay AD onset, by conserving locomotion function. Moreover, SAA diet reduces *hAβ42* levels at 14 days compared to YBD-fed flies (Fig2C), explaining the later activation of cell competition, as SAA-fed flies experience less disease until later in life. These findings align with evidence indicating that diets rich in fruits, vegetables, fish, legumes, and unsaturated oils, can reduce Aβ levels in humans (as reviewed in Díaz et al. 2022). In AD brains, IIS dysregulations cause insulin resistance and decrease insulin-degrading enzyme (IDE), which also degrades Aβ, thus promoting Aβ accumulation and plaque formation (Bedse et al. 2015). It is expected that diets that alleviate the burden of insulin resistance will promote Aβ clearance delaying AD progression. We speculate that SAA is a more balanced diet than YBD and may reduce the Aβ similarly, as IDE is conserved in *Drosophila* and mitigate Aβ neurotoxicity (Tsuda et al. 2010). Our results showed that diet modulates the elimination of unfit cells in AD and that locomotion correlates with cell competition, which in turn can be modulated by diet. Nevertheless, we cannot exclude that locomotion phenotypes may be partially due GMR to drive *GFP* and *hAβ42* expression, since is a driver known to induce toxicity and retinal degeneration (Escobedo et al. 2021; Kramer and Staveley 2003). As Currier *et al*. reviewed, flies’ behaviour relies heavily on the visual cues for several locomotion behaviours. Thus, flies will have some degree of locomotion decline induced by the nature of the driver used and regardless of disease or dietary effects (Currier, Pang, and Clandinin 2023). However, the fact that we were able to correlate locomotion restoration with cell competition activation in SAA-fed flies (Fig1L-L’’’), suggests that locomotion effects are related with cell competition and diet. Given that we used the same driver in both diets, we should observe the same degree of degeneration. Thus, the SAA diet might be sufficient to overcome GMR effect. Further studies are needed to understand how specific nutrients provided by diet and molecular signals relate to competition and their impact on ageing and neurodegenerative diseases. We believe our findings help bridge the gap in the field and shed light on new potential strategies for delaying AD progression through cell competition modulation with nutrition.

### Limitations of the study

Our work has contributed to understand hFWE function and conservation, and unravelled diet’s effects on the modulation of fitness fingerprints. However, there are some limitations. The undefined composition of YBD comprising yeast, syrups and flour, without a chemically defined composition, prevents identification of nutritional differences between YBD and SAA responsible for the effects seen. Manipulation of SAA diet components will allow us to overcome this limitation. For AD studies in *Drosophila*, we used Casas-Tinto model, where GMR drives expression of two copies of *hAβ42* (*GMR-Gal4, UAS-2xhAβ42*) (Casas-Tinto et al. 2011) and established a control model with one *GFP* copy driven by GMR (*GMR-Gal4, UAS-GFP*). Differences in the UAS count, construct insertion sites, and the recombination points between the GMR and UAS (despite being in the same chromosome), may affect the results, limiting direct comparisons between HEALTHY and AD flies. Therefore, data analysis was conducted within each genotype. Gene-Switch technology (Robles-Murguia et al. 2019) could address this constrain in future studies.

The locomotion assay requires careful interpretation, as GMR domain and retina manipulation may not be sufficient to induce motor dysfunctions and effects might stem from visual system dysfunction per *se*. However, locomotion effects align with cellular competition results. Future studies could use ELAV to manipulate all neurons and evaluate other behaviours, such as climbing activity. Phototactic response variability in the Buridan assay could also impact results, as less phototactic flies show reduced activity, preventing identification of effects (Colomb et al. 2012; Han et al. 2021). Furthermore, *hFWE* isoforms were expressed throughout development and compensatory effects may influence cellular and behavioural findings. Whether *Drosophila* Flower isoforms are regulated by *hFWE* overexpression remains unclear. Despite these limitations, our research provides valuable insights into *hFWE* functional conservation and diet-driven cell competition modulation in HEALTHY and AD flies, demonstrating that efficient unfit cell elimination enhances locomotion in disease and aging models.

## Methods

### Drosophila Maintenance

*Drosophila* melanogaster stocks were maintained at 25°C, with 60% humidity and a 12 h light/12 h dark cycle. Flies were kept in vials with a yeast-based diet (YBD Recipe) containing for 1L: 80g of Barley Malt Syrup (Próvida), 22 g of Beet syrup (Grafschafter), 80g corn flour, 18g of Instant yeast (Lesaffre), 10g Soy flour (Centazzi), 8g of Agar (NZYtech), 12ml Nipagin 15% and 8ml Propionic acid > 99% (Acros).

### Experimental protocols Fly brain dissection

For optic lobes, fly brains were dissected in cold PBS 1x, fixed in 3,7% PFA for 30 min at RT, washed with PBS-Triton-X 0.4% for 30 min and incubated with primary antibody (mouse anti-β-Amyloid, 17-24 [4G8] #SIG-39220 1:100 in PBS-T 0.4% and 5% of Normal Donkey serum (Jackson ImmunoResearch #017-000-121) overnight at 4ºC. After removing the primary antibody, the tissue was washed for 30 minutes with PBS-T 0.4%, and incubated with a secondary antibody (Alexa 488 #A-21202 Invitrogen 1:1000 in PBS-T 0.4%) at 4ºC overnight. The following day, the secondary antibody was rinsed, and the sample was washed with PBS 1x for 30 minutes. The optic lobes were mounted in Vectashield with DAPI (Vector Labs Inc #H-1200) on microscope slides and stored at 4ºC. Adult brains were mounted with a spacer to avoid compression of optic lobes. Samples were imaged on an 880 Zeiss confocal microscope.

### TUNEL staining

TUNEL kit assay (Roche cat# 03333566001) was used with some alterations compared to the manufacturer’s protocol. After brain fixation, brains were washed for 1 hour in 0.4% PBST and incubated with TdT buffer for 1 hour, both at room temperature. Afterwards, samples were incubated in TUNEL solution containing Terminal Transferase enzyme (3ul/ml) and biotin-16-dUTPs (Roche # 11093070910) (2ul/ml) diluted in TdT buffer (3ul/ml) for 2 hours at 37ºC. Reaction was stopped with STOP citrate buffer for 15 minutes at room temperature and then washed in PBS 1x for a total of 4-5 hours, with the PBS 1x being replaced every 45 minutes. Samples were then incubated with Streptavidin 488 or 647 (Invitrogen #S11223, #AB_2336066) diluted in PBS-T) with 10% Normal Donkey Serum (Jackson ImmunoResearch #017-000-121) at 4ºC overnight. The following day, the samples were washed for 2 hours with PBS 1x, replacing the PBS every 30 minutes.

### Diets protocol

Crosses were set up in a Vienna diet at a ratio of 3 females to 1 male per vial at 25ºC. After 3 days of egg laying, flies were discarded, and progeny allowed to develop and hatch. On the third day of hatching, F1 female flies were sorted according to the correct genotype and placed either in YBD or synthetic diets for 3, 14, 21, and 28 days. At these time points, we performed locomotion assays followed by dissection of fly brains. In the case of the synthetic diet, flies were fed with the FLYAA (referred as standard synthetic diet SAA in this study) developed by Piper et al. (2017).

### Locomotion assay – Buridan’s Paradigm

Buridan’s paradigm was used to evaluate the locomotion behaviour and performed as described (Colomb et al. 2012).Trajectories were analysed using Centroid Trajectory Analysis (CeTrAn) software (Colomb et al. 2012, (https://github.com/jcolomb/CeTrAn/releases/tag/v.4) providing eleven metrics to evaluate locomotion. Statistical analysis was performed using GraphPad software.

Mated females were collected on the second day of hatching for the behavioural experiments. Their wings were clipped to one-third of their length to prevent them from escaping the arena. The flies were then distributed one per vial and allowed to recover for 24 hours. One hour before the assay, flies were transferred to empty vials to prevent grooming behaviour in the arena. Flies were placed individually in the centre of the arena, and their locomotion behaviour was recorded for 5 minutes using the Buritrack software(Colomb et al. 2012). The assay was restarted if the flies jumped in a maximum of 2 times. Flies immobile for more than 1 minute were excluded from the assay. After each run, the flies were dissected. Locomotion assessment was performed 3, 14, 21 and 28 days of feeding. Flies were flipped every three days to ensure the nutritional quality of the diet.

### Image quantification

We have developed a new quantification method using Ilastik and MATLAB. Initially Ilastik, a machine learning tool for image classification, segmentation, and analysis, was used to identify pixels based on colour/intensity, edge and texture (Berg et al. 2019). A representative image is uploaded, and the user annotates the signal(s) of interest (TUNEL and hAβ42 – green and *azot*-red) and background iteratively, with real-time feedback on segmentation quality. Once satisfied with the network’s performance, the entire image batch is processed for segmentation. Ilastik generates probability masks indicating pixel signal. A custom MATLAB script then quantifies signals by loading a merged confocal image of z-projections with signals of interest. After drawing the region of interest (ROI), which is the optic lobe, the probability mask is loaded, and the size (for TUNEL - 17, for hAβ42 18 and *azot* 17) and intensity thresholds (0.05 for green signal and 0.1 for *azot*) are applied to eliminate noise and low-probability objects. The image is then binarised. The same size and threshold values were used for a given signal across all samples. After processing all images, another MATLAB script creates an Excel file with data from the .mat files. The data obtained is transferred to GraphPad graphical representation and statistical analysis.

### Statistical analysis

Our data do not assume a Gaussian distribution, and results are shown as median ± 95 % CI. Multiple comparisons were performed, applying the Kruskal Wallis test with the post-hoc Dunn’s test. Statistical analyses were performed with GraphPad Prism software. Results were considered significant at * p <0.05; ** p <0.01, *** p <0.001, **** p <0.0001.

## Author Contributions statement

C. C-R., C.B-P and E.M. designed the experiments, C. C-R. conducted the experiments and analysed data; C. C-R and J.J. developed the data analysis strategy, J.J wrote the MATLAB code; C. C-R., C.B-P and J.R.J wrote the manuscript, J.C. contributed to the flies’ dissections.

## Acknowledgments

We sincerely thank Sergio Casas-Tinto and Soraia Caetano for their kind assistance in revising the manuscript. Zita Carvalho-Santos for discussions, advice and support in several stages of this work. Additionally, Carlos Ribeiro for discussions and providing the reagents necessary, also his lab members Célia Baltazar and Inês de Hann Vicente for all the help preparing for the synthetic diet. Claúdia Almeida for discussions and Takashi Koyama for stocks. Lastly, we want to thank Champalimaud foundation fly, glass wash and ABBE platforms for all the background work that contributed to the development of this project. The Vienna Drosophila Resource Center, the Bloomington Stock Center, for sending stocks.

## Funding

Work in our laboratory was funded by Fundação D. Anna de Sommer Champalimaud e Dr. Carlos Montez Champalimaud, the European Research Council [Consolidator Grant to E.M.: Active Mechanisms of Cell Selection: From Cell Competition to Cell Fitness, 2014-2019; grant agreement ID 614964], the Portuguese Foundation for Science and Technology-FCT (PTDC/BIA-CEL/3594/2020 - DOI 10.54499/PTDC/BIA-CEL/3594/2020). C.C.-R. work was funded by FCT (SFRH/BD/137397/2018 and COVID/BD/153227/2023); J.R.J. work was funded SFRH/BPD/109659/2015, European Research Council Consolidator Grant #866237, Simons-Emory International Motor Control Consortium (Simons Foundation #717106); J.C work was funded by ERC Active Mechanisms of Cell Selection: From Cell Competition to Cell Fitness, 2014-2019; grant agreement ID 614964;C.B-P. work was funded by FCT; Fly platform was funded by CONGENTO LISBOA-01-0145-FEDER-022170, co-financed by FCT (Portugal) and Lisboa2020, under the PORTUGAL2020 agreement (European Regional Development Fund).

## Competing Interests

The authors declare no competing or financial interests.

## Data and Resource availability

All relevant data and details of resources can be found within the article and its supplementary information. The code generated for quantifications is available upon request.

## Notes

### Competing Interest Statement

The authors have declared no competing interest.

