## Supplemental Figures for "Dietary Interventions Modulate Cell Competition and Locomotor Decline in a Alzheimer’s Disease *Drosophila* Model"

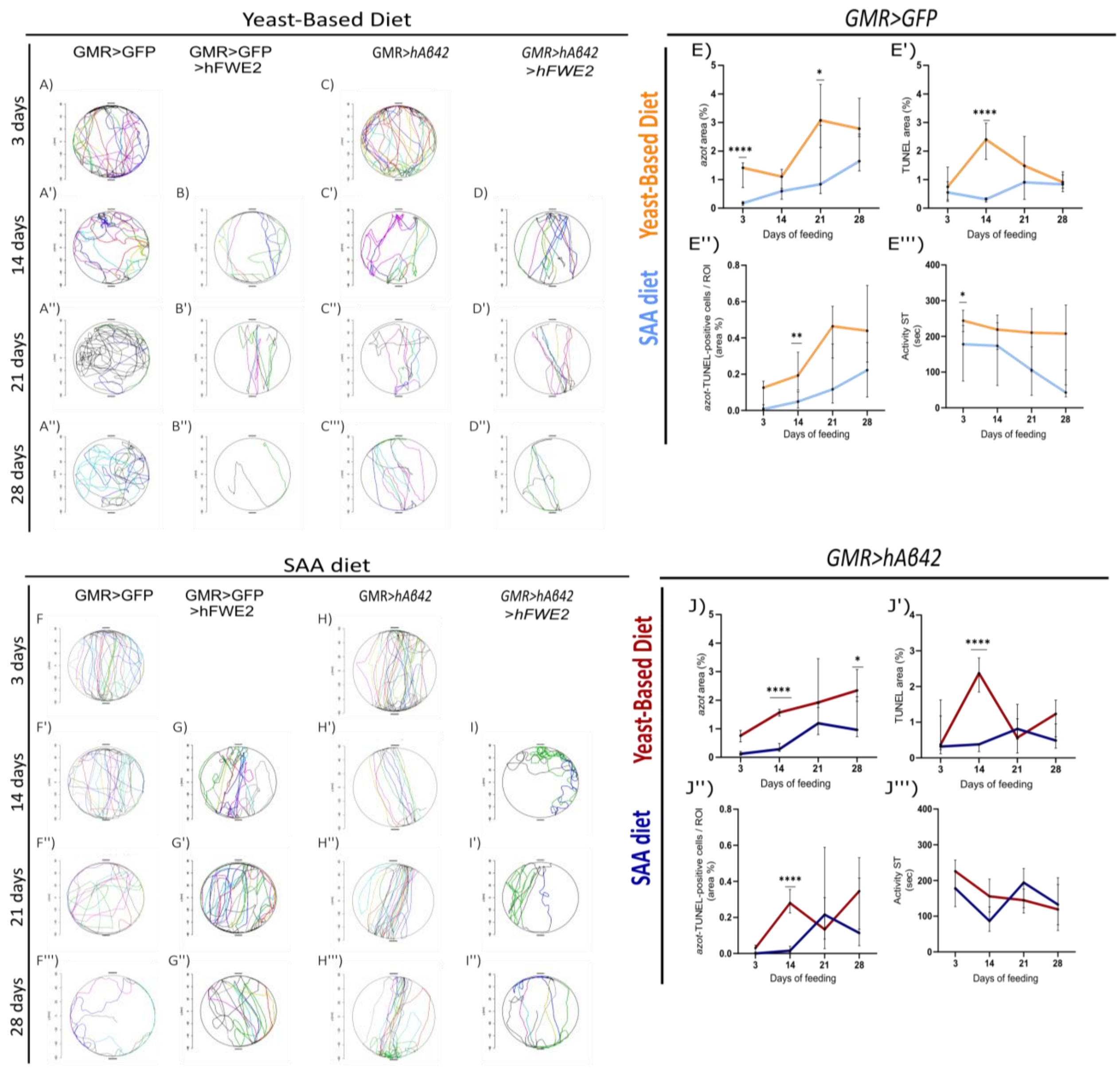


**Figure S1 - Effects of Yeast-based diet compared to the SAA diet –** (A-D'') Locomotion data of flies fed with YBD. (F-I'') Locomotion data of flies fed with SAA diet**.** (E-E''') Data from WT flies fed with YBD (orange) and SAA diet (light blue). (J-J''') Data from AD flies fed with YBD (red) and SAA diet (dark blue). (E,J) Area of *azot* normalized to the ROI (%). (E',J') Area of TUNEL normalized to the ROI (%). (E'',J'') azot-TUNEL-positive cells is the *azot* area colocalized with TUNEL area (%). (E''',J''') Fly's activity in seconds. All flies were fed for 3, 14, 21, and 28 days. ROI is the optic lobe. Data presented here is the same individually shown on Fig 1.

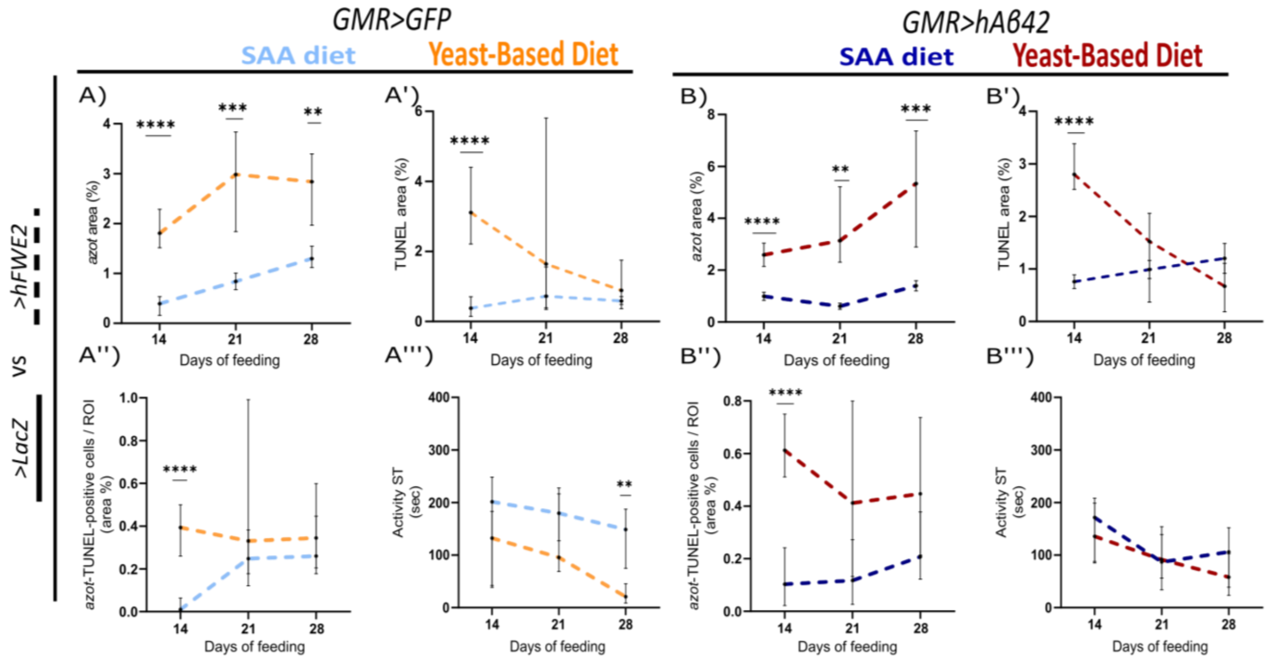


**Figure S2 – Comparison of hFWE2-expressing flies fed with SAA vs YBD.** (A-A''') WT flies expressing hFWE2 and fed with YBD (orange) and SAA diet (light blue). (B-B''') AD flies expressing hFWE2 and fed with YBD (red) and SAA diet (dark blue). (A, B) Area of azot normalized to ROI (%). (A',B') Area of TUNEL normalized to ROI (%). (A'',B'') azot-TUNEL-positive cells is the azot area colocalized with TUNEL area normalized to ROI (%). (A''',B''') Fly's activity in seconds. All flies were fed for 3, 14, 21, and 28 days. ROI is the optic lobe. Data presented here is the same individually shown on Fig 4 by the dashed lines.

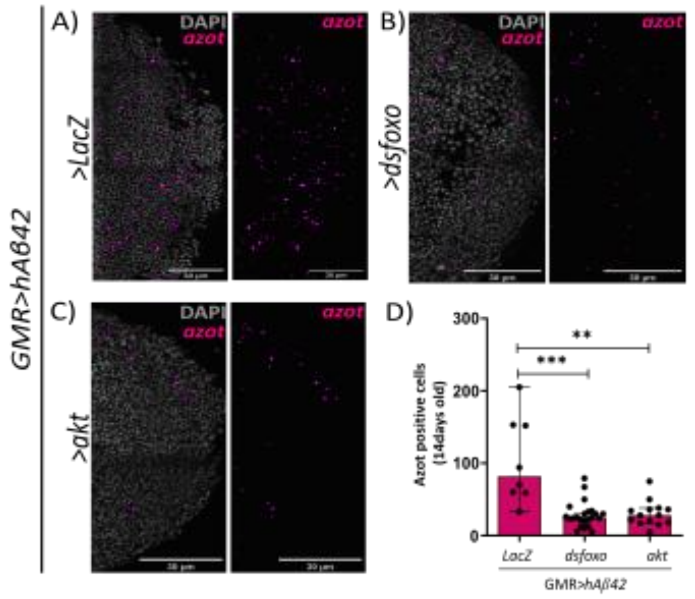


**Figure S3 –*azot* expression regulation by key metabolic regulators.** A) *azot* expression (in pink) when the control *LacZ* was expressed. In grey is the DAPI signal. B) *azot* expression when the *ds-foxo* was expressed. C) *azot* expression when the *UAS-akt* was expressed. D) Quantification of the *azot* levels seen in Figures A and B and C.
